# Detecting Direct Oral Anticoagulants in Trauma Patients using Liquid Chromatography-Mass Spectrometry: a Novel Approach to Medication Reconciliation

**DOI:** 10.1101/651075

**Authors:** Sudha Jayaraman, Jonathan H. DeAntonio, Stefan W. Leichtle, Jinfeng Han, Loren Liebrecht, Daniel Contaifer, Caroline Young, Christopher Chou, Julia Staschen, David Doan, Naren Gajenthra Kumar, Luke Wolfe, Tammy Nguyen, Gregory Chenault, Rahul J. Anand, Jonathan D. Bennett, Paula Ferrada, Stephanie Goldberg, Levi D. Procter, Edgar B. Rodas, Alan P. Rossi, James F. Whelan, V. Ramana Feeser, Michael J. Vitto, Beth Broering, Sarah Hobgood, Martin Mangino, Michel Aboutanos, Lorin Bachmann, Dayanjan S Wijesinghe

**Author notes:** **Corresponding Authors:** Sudha Jayaraman, MD MSc FACS, Associate Professor of Surgery, Division of Acute Care Surgery, Co-Director, Program for Global Surgery, Department of Surgery, VCU School of Medicine, VCU Health System; And Dayanjan S Wijesinghe, Assistant Professor, Department of Pharmacotherapy and Outcomes Sciences, VCU School of Pharmacy. **Disclosures:** The authors have nothing to disclose. Email addresses for all authors.

## Abstract

**Background:** Accurate medication reconciliation in trauma patients is essential but difficult. Currently there is no established clinical method of detecting direct oral anticoagulants (DOACs) in trauma patients. We hypothesized that a liquid chromatography-mass spectrometry (LCMS) based assay can be used to accurately detect DOACs in trauma patients upon hospital arrival.

**Methods:** Plasma samples were collected from 356 patients who provided informed consent including-10 healthy controls, 19 known positive or negative controls and 327 trauma patients over 65 years of age who were evaluated at our large, urban Level 1 Trauma Center. The assay methodology was developed in healthy and known controls to detect apixaban, rivaroxaban and dabigatran using LCMS and then applied to 327 samples from trauma patients. Standard medication reconciliation processes in the electronic medical record documenting DOAC usage was compared with LCMS results to determine overall accuracy, sensitivity, specificity and positive and negative predictive values (PPV, NPV) of the assay.

**Results:** Of 356 patients, 39 were on DOACs (10.96%): 21 were on Apixaban, 14 on rivaroxaban and 4 on dabigatran. The overall accuracy of the assay for detecting any DOAC was 98.60%, with a sensitivity of 94.87% and specificity of 99.06%, (PPV 92.50% and NPV 99.37%). The assay detected apixaban with a sensitivity of 90.48% and specificity of 99.11% (PPV 86.36% and NPV 99.40%). There were three false positive results and two false negative LCMS results for apixaban. Dabigatran and rivaroxaban were detected with 100% sensitivity and specificity.

**Conclusions:** This LCMS-based assay was highly accurate in detecting DOACs in trauma patients. Further studies need to confirm the clinical efficacy of this LCMS assay and its value for medication reconciliation in trauma patients.

**Study type:** diagnostic test

**Basic Science paper:** therefore does not require a level of evidence.

## Background

Medication reconciliation is universally recognized as an essential part of safe and reliable healthcare, and is particularly important in trauma patients.(1, 2) Trauma is the leading cause of death in the United States for those under 45 years old and cost over $600 billion per year.(3) However, the value of timely and accurate medication reconciliation in the trauma patient population remains underappreciated and its accuracy is dismal: 10-72%.(4-8) Trauma patients have unique problems which makes medication reconciliation more challenging – severe or distracting injuries, alterations in consciousness, intoxication, anxiety associated with trauma and the urgent nature of emergency care. The multiple transitions in care necessary for trauma care is also associated with an increased risk of medication errors.(9) These limitations make current medication reconciliation processes are inefficient, inaccurate and poorly suited for the trauma setting.(5, 10, 11)

Medication reconciliation has many steps, and therefore chances for error, since it involves obtaining and verifying a current medication list and clarifying that the medications and doses are appropriate before reconciling this list with the patient’s acute clinical treatment. It may not be possible to obtain information, such as current medication list, the name of pharmacy or primary care physician or time of last dose, from an incapacitated trauma patient. Verifying information may be difficult. Patients may use alternative sources of medications such as online pharmacies or have prescriptions from other health systems or cities. This information often cannot be rapidly or accurately obtained from incoherent or incapacitated trauma patients. All these factors will make it impossible to establish an accurate and complete list of medications to facilitate clinical decision making in the emergency setting.

Medication omissions, incomplete knowledge of medication usage and incorrect documentation from inaccurate medication reconciliation contribute to adverse drug events and medication errors.(5, 6, 12) This can be detrimental in a trauma patient on anticoagulation who does not receive reversal agents, or may have cardiac complications if beta-blocking agents are inadvertently withheld, or seizures if anti-seizure medications are not continued.(13, 14) Drug-drug interactions is an active area of research; common medications such as warfarin have over 200 identified drug interactions.(15) (16) The potential for these risks has led to medication reconciliation being considered as a National Patient Safety Goal by The Joint Commission accrediting all health facilities.(17)

A major concern, especially in older trauma patients, is the detection and reversal of direct oral anticoagulants (DOACs), such as apixaban, rivaroxaban and dabigatran, to combat acute hemorrhage. DOACs have been first line agents for atrial fibrillation and venous thromboembolism.(18, 19) Currently, no objective and widely available method exists to detect DOACs in the clinical setting.(20-22) Routine coagulation assays do not provide a complete assessment of the level of anticoagulation induced by DOACs.(23-25) Some tests such as thrombin time or Ecarin clotting time are feasible but not routinely available in the clinical setting.(26) Thromboelastograms do not reliably detect all DOACs. (27-29) The bleeding risk of DOACs has led to expedited FDA approval of reversal agents, but the of lack objective clinical tests to detect DOACs limits the use of these reversal agents.(30-32) Therefore, a better approach is needed to identify DOACs and facilitate accurate medication reconciliation in this high-risk patient population, to have good clinical outcomes.

Our group has studied the importance of medication reconciliation in trauma and potential applications from the field of metabolomics in the last few years.(4, 33) We hypothesized that a liquid chromatography-mass spectrometry (LCMS) based assay can accurately detect DOACs in trauma patients. LCMS-based technology has been increasingly used to study DOACs, develop methods of detection and determine clinical applications but has not yet been used in the trauma patient population.(34-37) We aimed to develop methodology for an LCMS-based assay to detect DOACs in healthy volunteers and patients with confirmed prescriptions for DOACs, apply it to a specimen bank of prospectively acquired samples from trauma patients evaluated at our large, urban Level 1 Trauma Center method, in a blinded fashion, and test its performance characteristics.

## Methods

### Patient Recruitment and Sample Collection

The Institutional Review Board at Virginia Commonwealth University approved this study. All subjects provided written informed consent prior to the study. A total of 356 subjects were enrolled: 327 trauma patients who presented acutely to the emergency department, 10 healthy volunteers, 10 outpatients who were on known doses of one of three DOACs of interest and 9 non-trauma emergency patients who were known to be either taking or not prescribed DOACs based on pharmacist confirmation (positive and negative controls). Healthy volunteers, defined as individuals who were not taking any medications, were recruited to donate fresh plasma samples for method development. Once consent was obtained, a standard blood draw procedure was performed on each subject. Drug free plasma was collected and then pooled, mixed, aliquoted and stored at −80 °C until data acquisition could be performed. Ten outpatients who were known to be prescribed DOACs were recruited from an outpatient cardiology clinic at our institution. Again, once informed consent was obtained, a standard blood draw procedure was performed on each subject and samples were again prepared and stored for analysis. At hospital arrival, routine laboratory samples are drawn from patients for standard clinical care. A pharmacist from the study team identified and recruited ten non-trauma emergency patients, based on convenience sampling, who were confirmed to be on one of the three DOACs of interest or none of them. After informed consent, residual plasma from their routine laboratory samples were obtained from the hospital lab and stored at −80 °C. All trauma patients who were 65 or older and presented to our Level 1 Trauma Center between July 2015 and March 2017 were eligible for this study. They were screened by clinicians and a study team member obtained consent from the patient or next of kin once they were confirmed to be eligible. Again, residual plasma from their routine laboratory samples were obtained from the hospital lab and stored at −80 °C.

A study-specific REDCap database was created by clinicians on the study team as a data collection tool and used to store clinical data from the electronic medical record and the institutional trauma registry, including current medication lists, comorbidities and clinical outcomes. Ten percent of the database records were independently audited using a random number generator to ensure accuracy of the stored data. The technical members of the study team were blinded to these clinical data during the LCMS analysis.

### Materials and Equipment

Dabigatran (D10090), dabigatran-d4 (D10092), rivaroxaban (R538000), rivaroxaban-d4 (R538002), apixaban (A726700) and apixaban-d3 (A726703) were purchased from Toronto Research Chemicals (North York, ON, Canada). The chemical structures of the DOACs and their internal standards are depicted in **Fig 1 Panel A**. LCMS grade water (W7SK-4), acetonitrile (A955-4) and ammonium formate (A1150) were purchased from Fisher Scientific (Waltham, MA, USA). The chromatographic separation column (ACQUITY UPLC BEH C18 Column, 130Å, 1.7 µm, 2.1 mm X 50 mm) was purchased from Waters (Milford, MA, USA). LCMS analyses were conducted using a Shimadzu Nexera UPLC system coupled to a Sciex QTRAP 6500 hybrid triple quadrupole linear ion trap tandem mass spectrometry system.

**Figure 1: Panel A.**
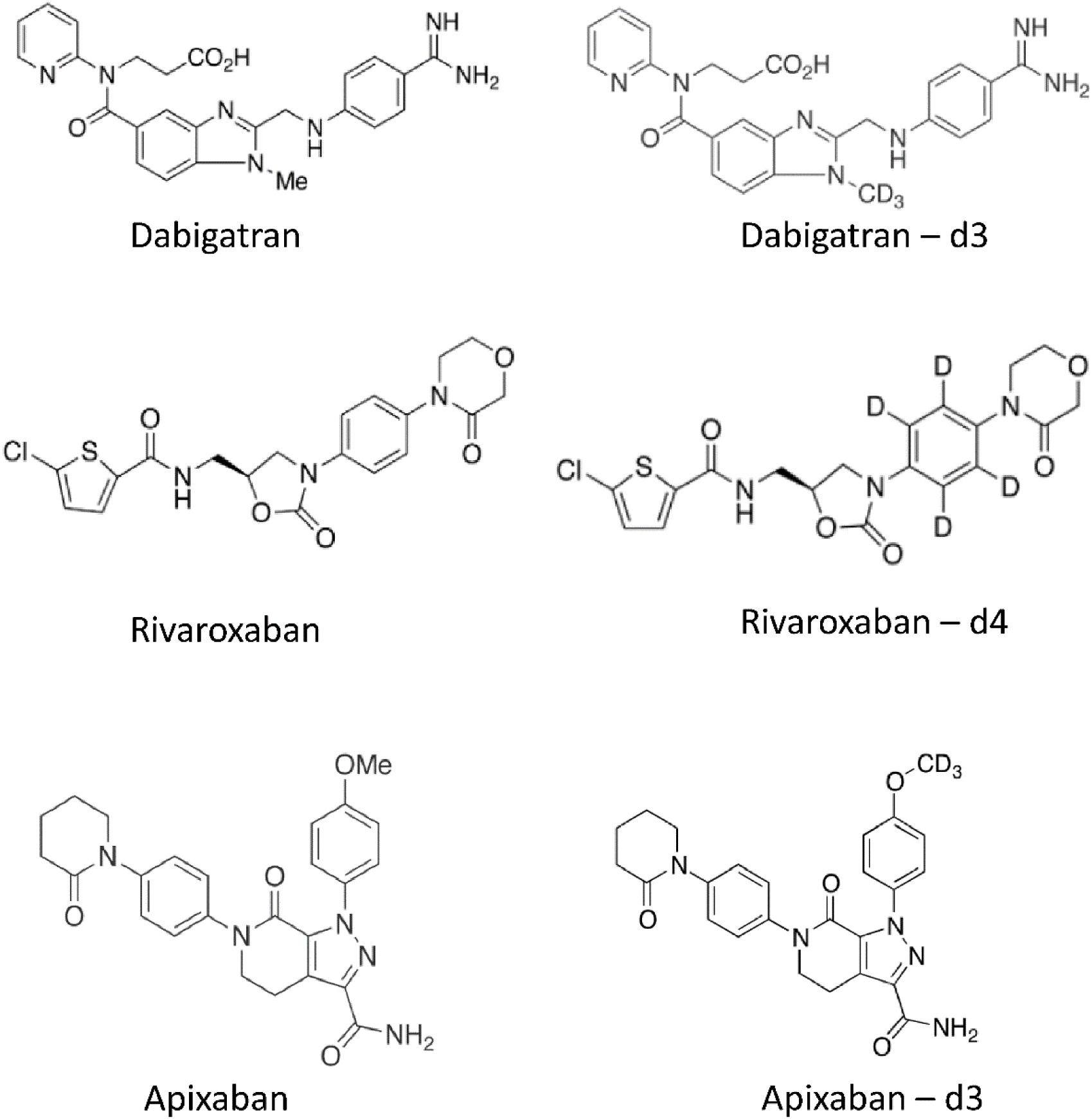

**Figure 1: Panel B.**
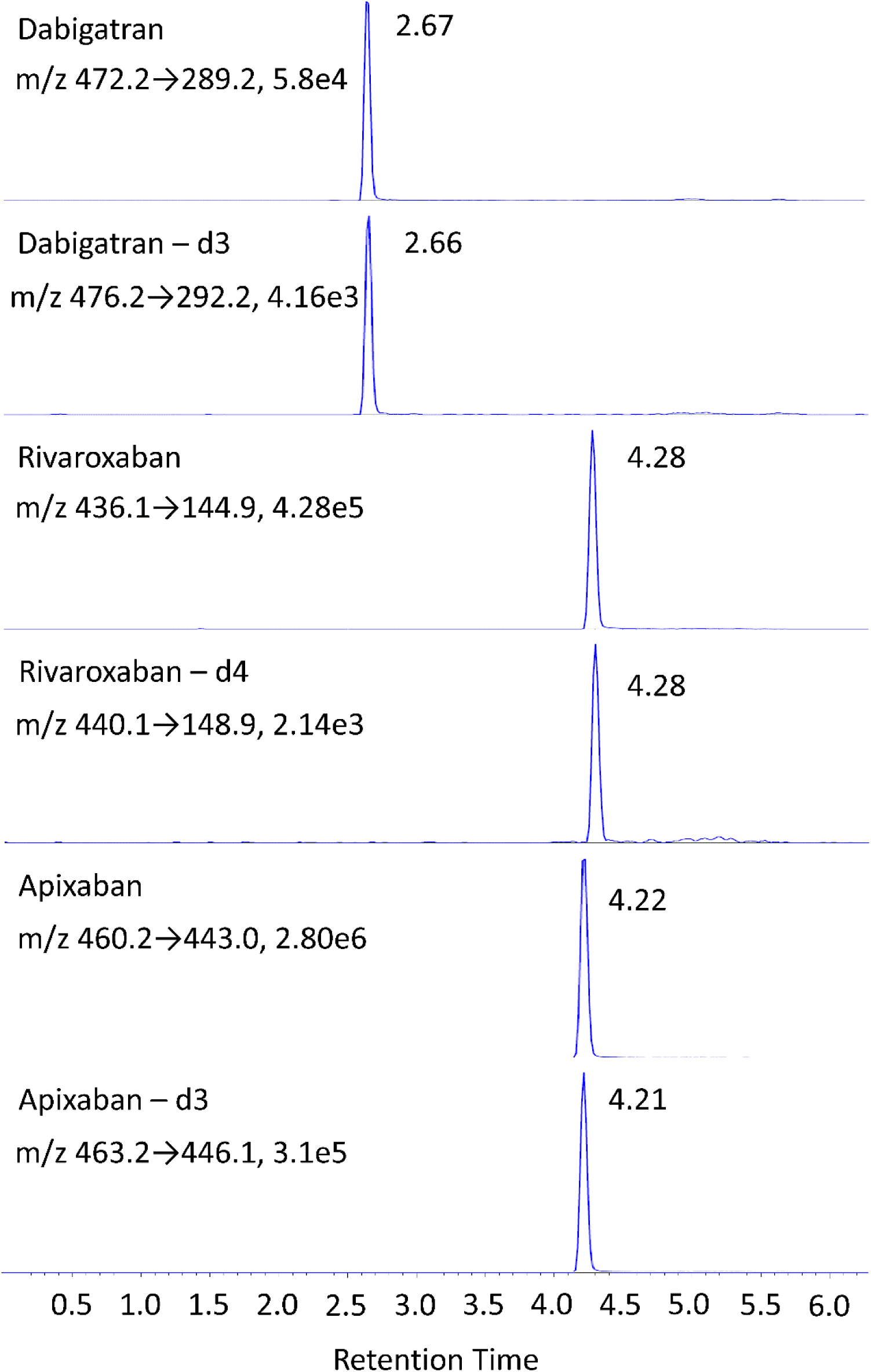

### Sample Preparation for LCMS Analysis

Calibrator (Cal) samples were prepared by spiking 50 µL each of drug free plasma with dabigatran, rivaroxaban and apixaban to obtain a 7-point calibration curve (10, 20, 80, 120, 240, 480 and 960 ng/mL). The same drugs were spiked into 50 µL of drug free plasma each to final concentrations of 80, 240 and 480 ng/mL to prepare quality control (QC) samples. Dabigatran-d3, Rivaroxaban-d4 and Apixaban-d3 internal standards were then spiked into the calibrators, QCs and patient samples to obtain a final concentration of 120 ng/mL. To allow rapid sample preparation, which would be necessary for implementation in a routine clinical laboratory, the sample pretreatment was limited to protein precipitation by dilution of each sample with acetonitrile at a ratio of 1:3 followed by the dilution of the supernatant at 1:4 with mobile phase A.

### LCMS Assay Parameters

An LCMS based assay was developed for the qualitative determination of the three direct oral anticoagulants of interest. Chromatographic separation parameters were established as follows: total flow rate was 400 µL/min. Oven temperature was set to 50°C. Injection volume was 10 µL. Autosampler cooler temperature was 15°C. Mobile phase A was water containing 2.5 mM ammonium formate titrated to a pH of 3.0. Mobile phase B was 100% acetonitrile. Analytes were separated using a 6.3 minute gradient based elution as follows: 0-0.7 minutes at 5% B. Linear increase of B to 30% by 2.8 minutes. B was held at 30% until 3.10 minutes and then increased linearly to 50% by 4.1 minutes and maintained at this level until 4.1 minutes. B was again increased linearly until 95% by 4.3 minutes and held until 4.9 minutes. B was then decreased to 15% by 5 minutes and held as such until 5.7 minutes. At 5.5 minutes, B was decreased to 5% and held at that level until 6.3 minutes to equilibrate for the next run. The analytes were monitored via multiple reaction monitoring (MRM) with the mass spectrometer operating in the positive mode. The monitored transitions were as follows: dabigatran (472.2→289.2), dabigatran-d3 (476.2→292.2), rivaroxaban (436.1→144.9), rivaroxaban-d4 (440.1→148.9), apixaban (460.2→443.0), apixaban-d3 (463.2→446.1). Dwell times for all analytes were held at 150 milliseconds. Instrument parameters for the analysis were: curtain gas at 30, source temperature at 300°C, nebulizer gas (gas1) at 40, drying gas (gas 2) at 60, ion spray voltage at 4500, declustering potential at 80, entrance potential at 10, CAD gas (nitrogen) at Low, collision energy at 40 and collision cell exit potential at 13. All samples, including positive and negative controls, were analyzed in a random and blinded fashion.

## Results

### LCMS Detection of DOACs in Plasma

The LCMS assay was developed by spiking DOACs into drug free plasma from ten healthy volunteers. Peaks were detected at the expected retention times for the appropriate mass transitions (**Fig.1 Panel B**). Samples from 10 outpatients known to be on treatment doses of DOACs were then tested in order to verify identification by the assay in a blinded fashion (**Table 1**). The assay accurately identified all except one as DOAC positive. A record review of this patient by the clinical team revealed that the patient had taken the medication 25 minutes prior to sample collection. Therefore, the plasma level of the medication had likely not reached therapeutic levels and did not cross the established detection threshold for the assay.

**Table 1:** LCMS Method Accuracy Assessment Using Positive Controls.

| **Patient** | **DOAC** | **Time of DOAC Dose (hr:min)** | **Time of Blood Collection (hr:min)** | **Detected (Y/N)** |
| --- | --- | --- | --- | --- |
| **1** | Rivaroxaban | 0830 | 1300 | Y |
| **2** | Apixaban | 1130 | 1400 | Y |
| **3** | Rivaroxaban | 1800- Prior Evening | 1410 | Y |
| **4** | Apixaban | 0900 | 1455 | Y |
| **5** | Rivaroxaban | 1900- Prior Evening | 1530 | Y |
| **6** | Apixaban | 1200 | 1545 | Y |
| **7** | Apixaban | 0900 | 1555 | Y |
| **8 \*** | Apixaban | 1620 | 1645 | N |
| **9** | Apixaban | 1000 | 1655 | Y |
| **10** | Dabigatran | 0600 | 1700 | Y |
| * Dosing was 25 minutes (1620) prior to blood collection (1645) |  |  |  |  |

### LCMS Assay Results Compared to Standard Medication Reconciliation in Trauma Patients

Once the assay methodology was established, all 356 samples were analyzed in a random fashion. The LCMS results were then compared to clinical data to assess the accuracy of the assay. Based on clinical data 39 of 356 patients were on DOACs (10.96%): 21 on apixaban, 14 on rivaroxaban and 4 on dabigatran (Table 2). When the LCMS-based assay results were compared against the current standard for medication reconciliation, the overall assay sensitivity and specificity for all DOACs were 94.87% and 99.06%, respectively. Overall positive predictive value (PPV) and negative predictive value (NPV) of the LCMS assay for the three DOACs were 92.50% and 99.37%, respectively. This represented an overall diagnostic accuracy of 98.60% for the assay to detect presence of any of the three DOACs.

**Table 2:**
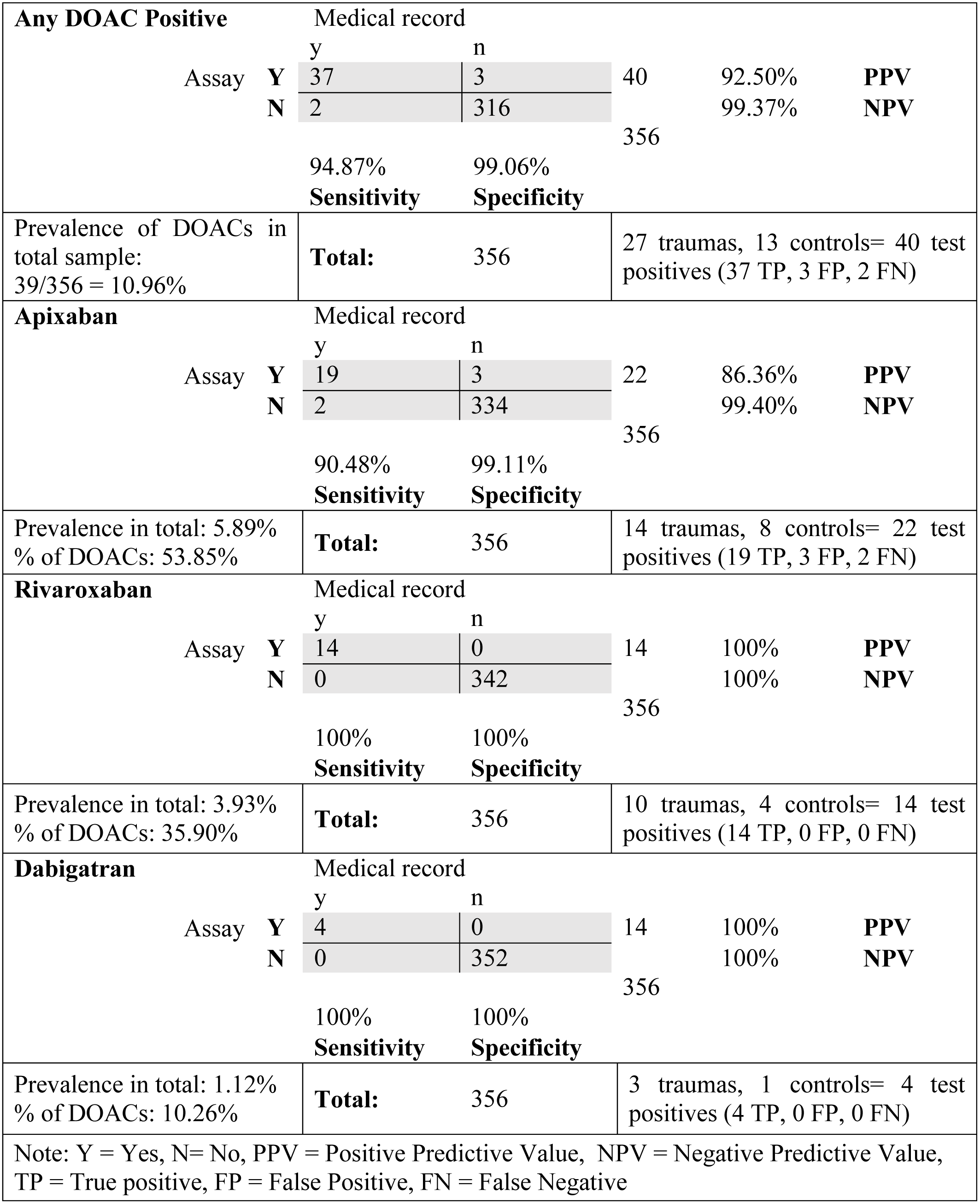
2×2 table for LCMS results for all DOACs and by each Individual DOAC.

For rivaroxaban and dabigatran, the appropriate drug was detected in all patient samples where the drug was recorded in the medical record and was not detected in samples from patients not known to be on these medications. Therefore rivaroxaban and dabigatran exhibited a 100% PPV and NPV in this study. Sensitivity and specificity for detecting apixaban were 90.48% and 99.11%, respectively and PPV and NPV were 86.36% and 99.40%, respectively. There were three LCMS results which were positive in patients with no record of apixaban therapy and two which were negative in patients with apixaban therapy noted in their medical records. (**Table 2**). All controls yielded the expected results.

## Discussion

Our study demonstrates that an LCMS-based assay can accurately detect DOACs in the trauma patient population where the current medication reconciliation processes are limited. The overall accuracy of the LCMS assay for detecting any DOAC was very high at 98.60% with approximately 95% sensitivity and 99% specificity. The assay accurately detected both rivaroxaban and dabigatran but the sensitivity and specificity of the most commonly used DOAC in our sample, apixaban, was lower (90.48% and 99.11%, respectively). This is still far superior to the 10-72% accuracy reported for medication reconciliation in the literature, but the false positive and false negative results need to be reviewed.

LCMS is highly precise and accurate and is routinely used to detect chemicals in other fields. (38-41) However, in this study, the assay may not have detected DOACs in patients’ plasma for several reasons: if the time interval between taking the medication and the blood sample being drawn was longer then the medication’s therapeutic window, i.e. 2-4 hours for full therapeutic effect and half-life of 5-14 hours for the DOACs evaluated in this study, the LCMS result will be negative, even if a patient has been prescribed and was taking the medication as directed. This scenario is quite likely in our trauma population as critically ill patients are transferred to our Level 1 facility for further care hours after their injury and their plasma, on arrival to our institution, may not contain detectable levels of the medication. Second, a patient who was injured before being able to take their daily dose will have a prescription for the DOAC but no detectable plasma level of this medication. Lastly, nonadherence to medication therapy may also correctly explain negative LCMS results where the medical record might indicate prescription of a DOAC.

In reviewing the medical record for the two samples with negative LCMS results, one patient was transferred seven hours after injury from a community hospital while the other was brought directly from the scene. Neither chart notes time of last dose of apixaban so it is unclear if any of the above factors contributed to the negative LCMS result when the patients were thought to be on apixaban. Nevertheless, when DOACs are not detectable by LCMS, patients are unlikely to have clinically relevant plasma levels that put them at risk for bleeding. Therefore, the assay results would have been potentially more useful to clinicians than the information available from the standard medication reconciliation process.

The LCMS assay may also have detected DOACs not captured in the electronic medical record, again highlighting the limitations of the current medication reconciliation process for trauma patients. (10, 42-44) When we reviewed the three patients who tested positive for apixaban, one was a patient who was transferred with a traumatic brain injury, which has been shown in the literature to limit medication history. (8) She had a history of stroke, hypertension and abdominal aortic aneurysm repair but had reported only being on sertraline and a “blood pressure medication”, although based on her comorbidities, she could conceivably have been prescribed an anticoagulant. The second case was a patient who did not know his medications and had his routine care at another facility, but had a history of myocardial infarction and stent placement. Again, this patient could reasonably have been prescribed an anticoagulant but that information was unavailable to the clinical team. The third case had medical record documentation showing that he was on a “blood-thinner” pre-injury, possibly warfarin. The accuracy of this information was unclear as the patient was altered. The LCMS assay, therefore, may have identified trauma patients at increased risk of bleeding who would not have been identified as such through standard medication reconciliation processes.

The discrepancy between the LCMS assay results and the clinical data highlights both the need for a robust medication reconciliation method to detect DOACs in trauma patients, and confirms the limitations of the current medication reconciliation processes. DOACs have replaced warfarin as the standard of care for anticoagulation.(45) Bleeding related complications from DOAC use has led to expedited approval of reversal agents by the FDA. However, there is no reliable qualitative or quantitative assay to rapidly detect DOACs and support treatment decisions to use these reversal agents. Indiscriminate use will be expensive and may potentially risk prothrombotic complications. An accurate mechanism to detect DOACs in trauma patients will be both clinically and economically valuable. However, the current medication reconciliation processes, to determine use of DOACs or other critical medications in trauma patients, are inadequate. (4)

The Joint Commission has nationally mandated medication reconciliation but existing processes do not offer objective and reproducible data on medication use and tend to be complex and time-intensive.(17) In the US, 137 million emergency department visits occurred in 2017, according to the Center for Disease Control, with nearly a third due to trauma.(46) Demographic shifts result in an increasingly older trauma patient population with multiple comorbidities who carry risks of poor outcomes.(47) Minimizing complications from medication errors in this high-risk patient population is critical.

LCMS is widely used to detect chemicals in other fields but has not been applied in the clinical setting until recently.(38-41) Multiple studies have now established LCMS-based methods to detect DOACs, demonstrated potential to monitor therapeutic levels or set the reference method for coagulation assays used to detect DOACs.(34, 36, 48, 49) These studies suggest that LCMS based assays can be useful for clinical care and, specifically, can serve as an objective source of information for medication reconciliation. Qualitative LCMS-based assays could facilitate medication reconciliation in trauma by detecting a wide variety of high-risk and critical medications. Other high-risk patient populations such as emergency surgery, sepsis, stroke, and cardiac patients may also benefit from the clinical application of this technology, suggesting future directions for research. (50) Applications of LCMS technology have several major logistical challenges before they are likely to be widely available in the clinical setting. Mass spectrometry is available in many hospital clinical labs across the U.S., but a lack of hardware capability to move beyond batch-mode processing and the need for significant technical expertise to perform mass spectrometry limit the widespread use of LCMS-based assays, particularly in smaller hospital systems.(33) However, this technology is rapidly evolving and likely to be more widely available with time.

This study has several limitations. First, traditional EMR-based medication reconciliation is the “gold standard” for comparison and is in itself highly flawed. Therefore, “inaccurate” results obtained via LCMS might in fact have been correct, and simply reflected the flawed traditional medication reconciliation process. Second, while the overall sample size was large, the prevalence of DOACs in our study was only 11% and so larger sample sizes may be necessary to study this assay in more detail. Lastly, each individual DOAC represented a smaller proportion of the sample size and we did not include the most recently approved DOAC – edoxaban in our study.

In conclusion, we found that an LCMS-based assay was highly accurate in detecting DOACs and may represent a promising objective method of performing medication reconciliation in the trauma patient population. This novel application of this technology may be especially valuable where accurate and fast medication reconciliation is crucial, but difficult.

## Acknowledgements

Numerous nurses, pharmacists and clinical staff who attempt to perform medication reconciliation on all of our trauma patients.

## Author Contributions

Study design: SJ, JHD, DSW

Patient enrollment: SJ, JHD, JH, LL, TN, MBA, RJA, JDB, PF, SG, SWL, LDP, EBR, APR, JFW, VRF, MJV, BB

Data acquisition: SJ, JHD, JH, LL, DC, CY, CC, JS, DD, NGK, GC, DSW

Data analysis: SJ, JHD, LL, DSW, LW

Initial drafting of the manuscript: SJ, JHD, DSW

Critical revisions: SJ, JHD, SWL, JH, LL, DC, CY, CC, JS, LW, DD, NGK, GC, RJA, JDB, PF, SG, LDP, EBR, APR, JFW, VRF, MJV, BB, SH, MM, MBA, LB, DSW

